# Comparative analysis of single nucleotide polymorphisms and microsatellite markers for parentage verification and discovery within the equine Thoroughbred breed

**DOI:** 10.1101/2021.07.28.453868

**Authors:** P. Flynn, R. Morrin-O’Donnell, R. Weld, L. M. Gargan, J. Carlsson, S. Daly, H. Suren, P. Siddavatam, K. R. Gujjula

## Abstract

Short tandem repeat (STR), also known as microsatellite markers are currently used for genetic parentage verification within equine. Transitioning from STR to single nucleotide polymorphism (SNP) markers to perform equine parentage verification is now a potentially feasible prospect and a key area requiring evaluation is parentage testing accuracies when using SNP based methods, in comparison to STRs. To investigate, we utilised a targeted equine genotyping by sequencing (GBS) panel of 562 SNPs to SNP genotype 309 Thoroughbred horses - inclusive of 55 previously parentage verified offspring. Availability of STR profiles for all 309 horses, enabled comparison of parentage accuracies between SNP and STR panels. An average sample call rate of 97.2% was initially observed, and subsequent removal of underperforming SNPs realised a pruned final panel of 516 SNPs. Simulated trio and partial parentage scenarios were tested across 12-STR, 16-STR, 147-SNP and 516-SNP panels. False-positives (i.e. expected to fail parentage, but pass) ranged from 0% for 147-SNP and 516-SNP panels to 0.003% when using 12-STRs within trio parentage scenarios, and 0% for 516-SNPs to 1.6% for 12-STRs within partial parentage scenarios. Our study leverages targeted GBS methods to generate low-density equine SNP profiles and demonstrates the value of SNP based equine parentage analysis in comparison to STRs - particularly when performing partial parentage discovery.

## Introduction

A core biological question often asked, is to who the parents of an offspring in question are and such analysis is commonly termed as parentage verification. Typical methods employed to perform parentage verification utilise molecular technology to analyse simple Mendelian inheritance principles between offspring, paternal and maternal candidates (Jones *et al*. 2010). This concept of molecular based parentage verification is currently applied within a range of diploid species including the equine species and serves as a key tool to ensure integrity of pedigree recording, when testing offspring against nominated sires and dams (Marklund *et al*. 1994, Binns *et al*. 1995, ISAG 2012).

Early equine parentage verification techniques utilised antigen-based blood typing markers (Bowling *et al*. 1985) and advancements in molecular technology facilitated a feasible change to using STR markers, which enhanced parentage verification accuracy and process efficiencies in comparison to blood typing methods (Bowling *et al*. 1997). Typical equine STR profiles consist of a minimum 12 markers recommended by The International Society of Animal Genetics (ISAG 2012) and STRs continue to serve as a robust and accurate choice of marker when performing parentage verification and population genetics studies across equine breeds (Lee *et al*. 2006, Tozaki *et al*. 2001, van de Goor *et al*. 2011 and Rahimi-Mianji *et al*. 2015). Several analogous species have similar industry-based parentage verification needs i.e. bovine and ovine (Heaton *et al*. 2002 and Heaton *et al*. 2014) and implementation of SNPs as a choice of molecular marker have proven astute, enabling respective industries to benefit from in-tandem SNP based applications such as: accurate parentage verification/discovery (Hill *et al*. 2008, Fernandez *et al*. 2013, McClure *et al*. 2015, Yu *et al*. 2015 and Berry *et al*. 2019), trait discovery (Nicholas *et al*. 2003), genomic relationship matrices (Makgahlela *et al*. 2013 and Moore *et al*. 2019), inbreeding monitoring (Forutan *et al*. 2018, Todd *et al*. 2018 and McGivney *et al*. 2020) and genomic trait selection (Wiggans *et al*. 2012 and Spelman *et al*. 2013). Consequently a transition from STR to SNP markers to perform SNP profiling for routine parentage verification needs, may be one of the first steppingstones to enable the wider equine industry leverage value adds of SNP based analysis applications, in tandem with meeting parentage testing requirements.

Specific down-stream application needs often drive selection of molecular marker type. SNPs have proven to be a robust marker choice across various species when performing parentage verification (Flanagan *et al*. 2019), with choice driven by attributes such as simple biallelic variation, multiplexing capabilities (ranging 10’s to millions of SNPs per individual), lower mutation rates (e.g. SNPs ∼10^−8^ vs STRs ∼10^−3^) (Butler *et al*. 2007) and cost-effective genotyping technologies (De Donato *et al*. 2013 and Flanagan *et al*. 2019). One such genotyping technology that has been successfully used to support parentage verification genotyping needs (Whalen *et al*. 2019), is genotyping by sequencing (GBS). One specific use of GBS enables effective targeted SNP multiplex genotyping (Longeri *et al*. 2019) – lending GBS as a candidate genotyping method to support generation of low-density equine SNP profiles for use within parentage analysis.

Global standardisation of molecular marker profiles and their use within animal parentage verification is currently governed by various institutes, inclusive of ISAG (https://www.isag.us/) and the currently bovine focused International Committee of Animal Recording (ICAR - https://www.icar.org/). Specifically, within the equine Thoroughbred breed - the International Stud Book Committee (ISBC - https://www.internationalstudbook.com/) governs acceptance criteria for Thoroughbred studbook entry and molecular based parentage verification is a core registration requirement (ISBC 2020). To date ISAG have performed three pilot equine SNP comparison tests (2017, 2019 and 2021 SNP horse comparison tests - SNP-HCT) which utilised a 154 pilot SNP panel (reduced to 147 SNPs for 2021 SNP-HCT: https://www.isag.us/Docs/ISAG_Horse_SNP_Panels_2020.xlsx) consisting of 101 “Etalon” SNPs (Holl *et al*. 2017) and 53 “Tozaki” SNPs (Hirota *et al*. 2010). Assessment of performance across laboratory and genotype platforms is currently ongoing for both Etalon and Tozaki SNP panels, to establish an official recommended ISAG SNP parentage panel of robust and informative nature (https://www.isag.us/Docs/EquineGenParentage2017.pdf).

A core objective of this study was to establish a GBS based equine SNP panel, including current ISAG pilot SNPs and an extra SNP panel to facilitate both parentage verification (i.e. against nominated parents) and accurate sire and dam discovery (i.e. assigning parents against a reference database, without any prior parental details). A subsequent objective was to estimate the effectiveness of this SNP panel in comparison to STRs, when performing parentage verification and sire/dam discovery within the closed Thoroughbred equine breed. Accumulatively these objectives aim to assist the global equine community within an ongoing decision-making process whether to officially recommend SNPs as an approved molecular marker type, when performing equine parentage analysis.

## Materials and methods

### Sample selection

A random selection of n=349 anonymous Thoroughbred horses, which previously underwent STR based parentage verification (performed at Weatherbys Scientific), were used to perform additional SNP based parentage verification testing. Initially 40 out of a total 349 horse samples were used to establish an extra 429 SNP panel for addition to the ISAG pilot 154 SNP panel. The remaining 309 out of 349 horse samples were used to perform simulated parentage testing (PT). The 309 samples consisted of 157 Trio-Case samples (N=55 offspring, N=47 sires and N=55 dams) which were previously STR parentage verified against nominated parents. A further 152 samples (N=67 sires and N=85 dams), with no known parental relationship to 55 offspring, were also included to increase the scale of simulated parentage testing analysis. Furthermore N=20 DNA extracts from ISAG HCT 2019 equine pilot SNP comparison test (DNA prepared by ISAG 2019 duty laboratory: Eurofins Medigenomix, Germany) were included to allow for genotype performance and genotype concordance analysis (Table 1).

**Table 1:**
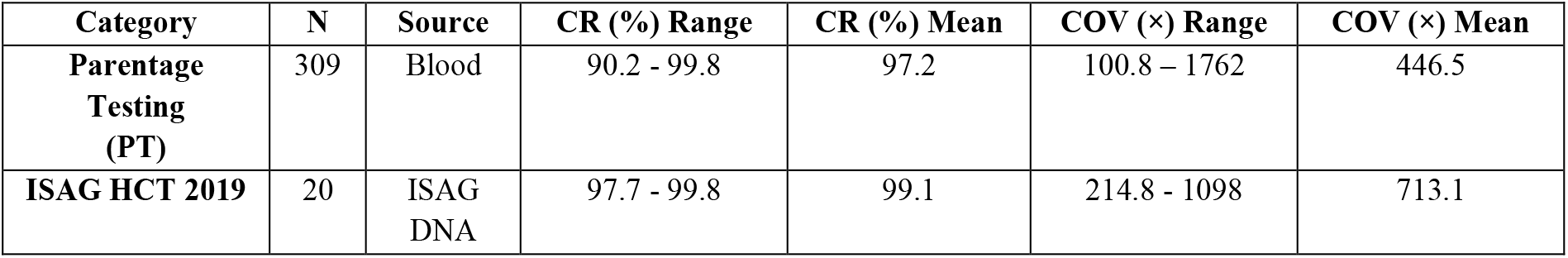
Sample numbers (N) and sources across category types, with sample call rate (CR) and sequence coverage (COV) based on overall 562 SNP panel.

| Category | N | Source | CR (%) Range | CR (%) Mean | COV (x) Range | COV (x) Mean |
| --- | --- | --- | --- | --- | --- | --- |
| Parentage Testing (PT) | 309 | Blood | 90.2 - 99.8 | 97.2 | 100.8 – 1762 | 446.5 |
| ISAG HCT 2019 | 20 | ISAG DNA | 97.7 - 99.8 | 99.1 | 214.8 - 1098 | 713.1 |

### GBS SNP marker selection and panel design

Using archived equine blood samples housed at Weatherbys Scientific, 40 out of 349 samples underwent DNA extraction using DNeasy Blood and Tissue Extraction Kit (Qiagen) and resulting DNA was subsequently genotyped on the Affymetrix 670K equine SNP array (Schaefer *et al*. 2017). Samples with genotype call rates >97% (i.e. all 40 samples) underwent SNP pruning starting from 670,796 SNPs to a panel of 429 SNPs (c.f. Fig. 1), of which core pruning considerations were to ensure selection of SNPs which genotyped well, were of informative nature and lastly - common to both Affymetrix 670K and Illumina 54K (McCue et al. 2012) equine SNP arrays, to potentially allow for backward compatibility to horses previously genotyped on either SNP arrays. All 101 Etalon and 5 out of 53 Tozaki SNPs were identified on the current Affymetrix 670K array and these 106 SNPs were isolated during SNP pruning to establish the extra 429 SNP panel. Furthermore, to allow for sex confirmation checks when performing parentage verification - 1 Y Chromosome SNP was selected from Affymetrix 670K array (AX-102953762) and included within GBS panel design.

**Fig. 1:**
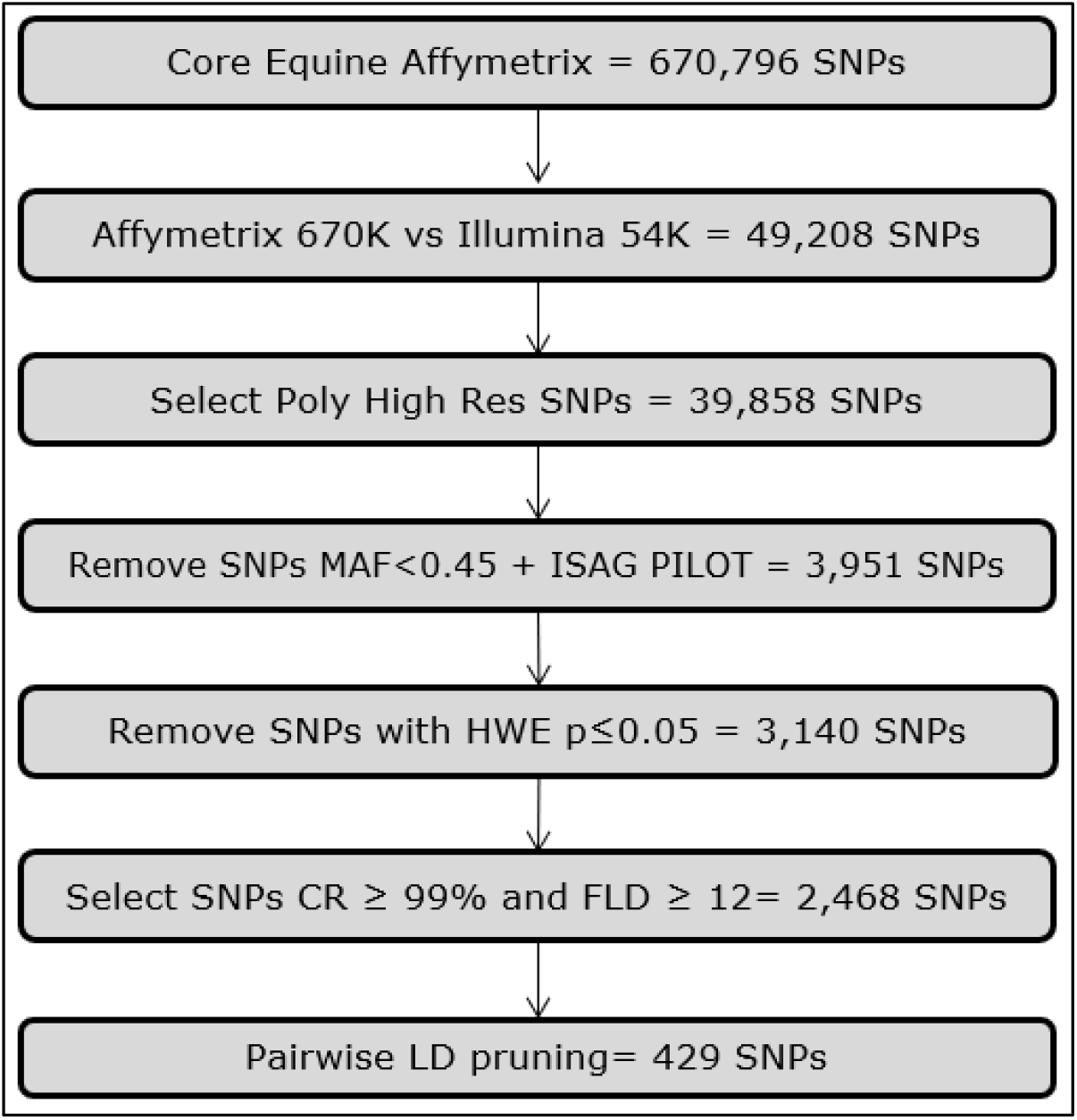
Range of SNP pruning criteria used to establish extra 429 SNP panel and associated SNP numbers post implementation of each criterion. Pruning criteria consisted of SNP selection as follows – common SNPs between Affymetrix 670K (Schaefer et al. 2017) and Illumina 54K (McCue *et al*. 2012) Equine SNP arrays (to allow for backward compatibility with previously SNP array profiled horses), selecting Polymorphic High Resolution SNPs (Affymetrix specific QC metric), removing SNPs with a minor allele frequency (MAF) <0.45, removing ISAG pilot SNPs, removing SNPs with a Hardy Weinberg Equilibrium (HWE) p≤0.05, selecting SNPs with Call Rate (CR) ≥ 99%, selecting SNPs with Fisher’s Linear Discriminant (FLD - Affymetrix specific QC metric) ≥ 12 and selecting SNPs post performing within-chromosome linkage disequilibrium (LD) pruning (indep-pairwise 50 5 0.04).

In total 584 SNPs consisting of - 101 Etalon, 53 Tozaki, 429 Extra and 1 Y-Chromosome SNP (Table S10) were submitted to the AgriBusiness bioinformatics team at Thermo Fisher Scientific for AgriSeq™ GBS panel design. To maximise GBS panel performance, all SNPs underwent re-mapping to the recent EquCab3.0 (Kalbfleisch *et al*. 2018) equine genome build where it was noted that flanking sequences for 13 SNPs (2 ISAG - BIEC911841 & BIEC2-462271 and 11 extra SNPs) mapped to multiple locations and as a result the 11 extra multi mapping SNPs were excluded; however both multi-mapping ISAG pilot SNPs were included due to their core panel origins. Post *in-silico* SNP removal, 573 SNPs remained for which chromosome, position and flanking SNP sequences were used for the GBS panel design and manufacturing (Table S1).

### GBS SNP panel performance, informativeness and concordance

Aliquots from all 309 PT horse blood samples (inclusive of 55 trio cases) underwent two rounds of cell lysis using HPLC grade water, whilst centrifuging and removing supernatant after each wash, along with a final clean up using Chelex-100 (20% solution) and Proteinase-K (20mg/ml) solutions. All 309 Chelex-based DNA extracts and 20 ISAG HCT 2019 DNA extracts, underwent genotyping using our synthisised equine genotyping by sequencing (EQ-GBS) SNP panel. GBS libraries were generated using the Applied Biosystems AgriSeq HTS Library Kit (Thermo Fisher Scientific). The amplified libraries were sequenced on Ion S5™ (Thermo Fisher Scientific), and genotype calls generated using Torrent Variant Caller (TVC) v5.12.2 (Thermo Fisher Scientific).

Genotype quality control metrics were applied at both a sample and SNP level. Dual sample level acceptance criteria of a genotype call rate ≥90% and a ≥100X average sequence coverage, along with a SNP level acceptance criteria of call rate ≥90% were applied. Genotype call thresholds within SNP were subject to a minimum TVC quality score of 10 and minimum coverage of 10X. To assess panel informativeness, minor allele frequencies (MAFs) were calculated using PLINK --freq (Purcell *et al*. 2007) for 254 PT (i.e. 309 PT samples with offspring removed) across ISAG pilot SNP, Extra SNP and overall EQ-GBS SNP (i.e. ISAG pilot and Extra SNPs) panels. *In-silico* MAFs were also calculated across 36 global-wide equine breed groups for Extra SNP panel via utilising multibreed Illumina 54K SNP data available from the study of Petersen *et al*. 2013.

Ten out of the twenty ISAG HCT 2019 samples were also genotyped in duplicate - allowing for “Within-Platform” reproducibility testing i.e. re-genotyping samples on the same platform. Furthermore 5 out of the 40 horse samples that underwent Affymetrix 670K Equine SNP genotyping to establish 429 Extra SNPs, were also genotyped using the EQ-GBS SNP panel - allowing for “Across-Platform” reproducibility testing i.e. Affymetrix 670K SNP Array versus Ion S5 GBS SNP panels.

### STR data generation

Historical STR genotype profiles, previously generated for STR based parentage verification needs, were available for all 309 PT samples. Individual profiles consisted of 17 equine STRs, inclusive of the minimum 12 recommended ISAG STRs (Table S2). As STR marker *LEX3* maps to Chr X, this marker was not included in parentage analysis simulations - leaving 16 autosomal STR genotype profiles per horse.

### SNP and STR parentage simulations

Simulated trio parentage testing was performed using all 309 PT samples, i.e. 55 offspring against 114 possible sires and 140 possible dams or alternatively 877,800 trio combinations i.e. offspring, sire and dam. Furthermore, 13,970 simulated partial combinations i.e. offspring, sire or dam were tested. Both trio and partial combination types included true “Expected to Pass (ETP)” parentage combinations for the core 55 offspring. All parentage combinations were tested using both SNP (Pilot ISAG and EQ-GBS) and STR (12 minimum recommended ISAG and all 16 markers) datasets. The number of mismatching markers per parentage combination were counted using bespoke PYTHON scripts (scripts for research use only and available at discretion of authors), which implemented Mendelian inheritance checks to calculate Mendelian error mismatch counts per pedigree combination across both STR and SNP marker types. A Pre-Analysis SNP run was performed to identify and remove SNPs with an above average percentage of Mendelian mismatch inconsistencies observed within ETP “offspring, sire and dam” parentage trio combinations for the core 55 offspring.

Subsequently two performance parentage testing measures (Strucken *et al*. 2016) were assessed: 1) Separation value and 2) False positive/negative case counts. 1) Separation value is defined as the difference between minimum number of mismatches for “Expected to Fail (ETF)” offspring-parents and maximum number of mismatches in ETP offspring-parents, scaled by dividing by number of markers. A greater than zero separation value indicates perfect separation is achievable and zero or negative separation values indicating non-perfect parentage test performance. 2) Exact counts of incorrectly failing (false-negative) and incorrectly passing (false-positive) trio/partial parentage combinations were also calculated as a proportion of false-negative and false-positive parentage combinations, with a ≥1% SNP and ≥1 marker STR mismatch exclusion threshold implemented when assigning parentage.

Average Non-Exclusion Probabilities were also estimated for categories One-Candidate-Parent (NE:1P) and Candidate-Parent-Pair (NE:PP) using Cervus 3.0.7 (Marshall *et al*. 1998 and Kalinowski *et al*. 2007). Estimates were performed using the 254 PT sample set (i.e. 309 PT samples with offspring removed) for all STR and SNP panel combinations. Probabilities of Exclusion (PE) were further calculated for both 1P and PP categories by subtracting respective NE values from 1, i.e. PE=(1-NE).

## Results

### GBS SNP panel

Post panel synthesis, 562 of 573 attempted SNPs were present on the synthesised EQ-GBS SNP panel i.e. a 98.1% design turn out. The 562 SNPs consisted of – 151 ISAG pilot (98 Etalon and 53 Tozaki), 410 Extra and 1 Y-Chromosome SNP(s) (Table S10). It was noted that 3 out of 154 ISAG pilot SNPs where not present in the final primer synthesis – BIEC2-476920, BIEC2-189666 and BIEC2-245923. The number of SNPs per autosomal chromosome ranged from 8 to 32, with a mean number of 18.5 SNPs per autosomal chromosome (Fig. 2). The average distance between the 562 SNPs across chromosomes was ∼3.95Mb and uniform genome coverage was observed as per Fig. S1.

**Fig. 2:**
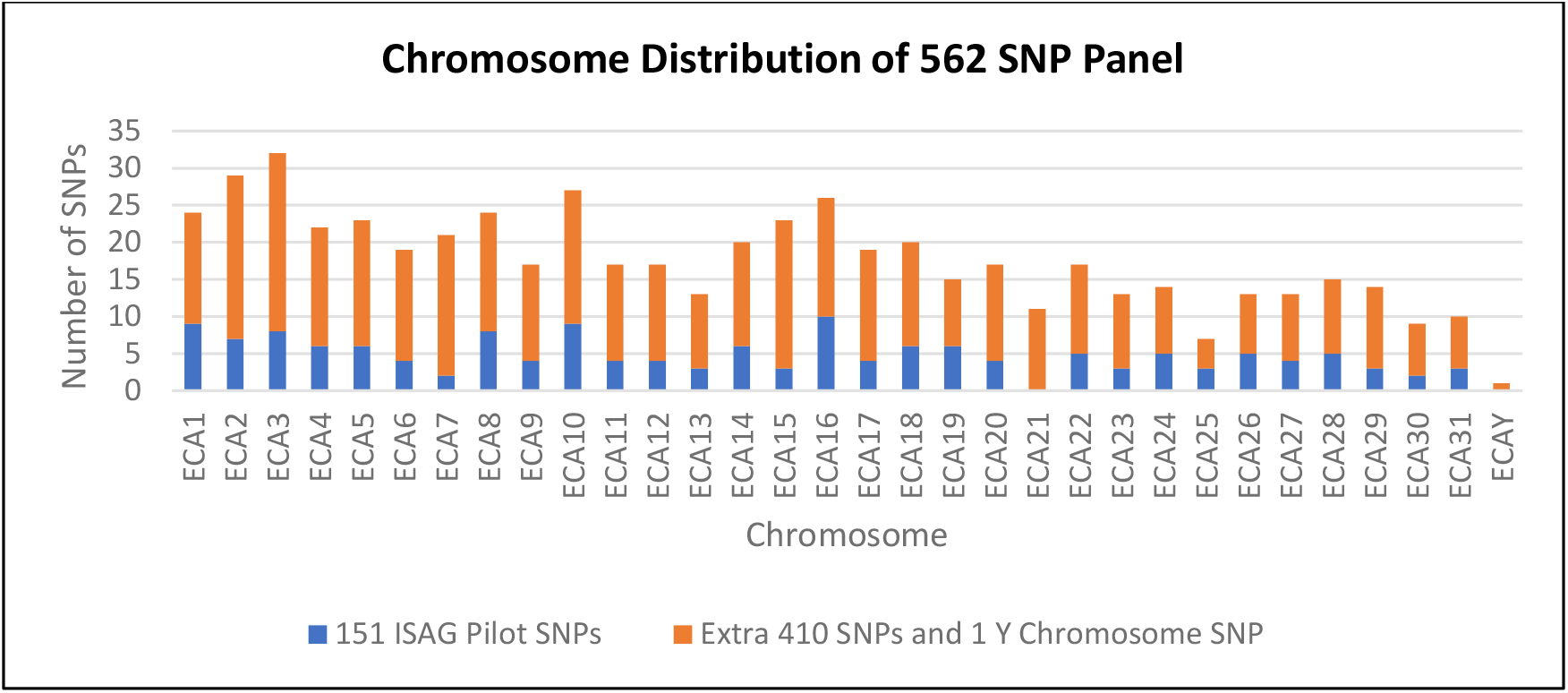
Distribution of 562 SNPs across all 31 autosomal and Y chromosomes (ECA), at a per chromosome level. Panel breakdown across both 151 ISAG pilot SNPs, Extra 410 SNPs and 1 Y Chromosome SNP shown in blue and orange respectively.

Sample call rates ranged from 90.2 - 99.8% (mean = 97.2%) for the 309 PT samples and 97.7% to 99.8% (mean = 99.1%) for 20 ISAG HCT 2019 samples (Table 1). SNP call rates were calculated across both PT and ISAG HCT 2019 samples with 523/562 and 548/562 SNPs having call rates ≥90%, within respective category types (Table 2 and Fig. S2). The panel of 523 SNPs (145 ISAG pilot and 378 Extra) with call rates ≥90% were used for further MAF, concordance and reproducibility testing. Minor allele frequencies were calculated using 254 PT samples across 145 ISAG pilot, 378 Extra and 523 EQ-GBS SNP panels. The 378 Extra SNP panel displayed the highest MAF of 0.40 across individuals, in comparison to 0.34 for the 145 ISAG pilot SNPs and combining both panels resulted in a MAF of 0.38 across 523 EQ-GBS SNPs (Table 2 and Fig. S3-S5). *In silico* multi-breed MAF values were based on an average of ∼371 (SD = 3.8) comparable Extra SNPs to Petersen *et al*. 2013 genotype data, and MAFs ranged from 0.25 (Mangalarga Paulista) to 0.41 (Paint and Quarter horse), with an across breed MAF average of 0.33 (Table S3 and Fig. S8).

**Table 2:** Number of SNPs from 562 panel within each sample type category, across a range of call rate (CR) categories. Minor allele frequencies (MAF’s) for ISAG pilot, Extra, and Combined EQ-GBS SNP panels as per PT samples.

| Sample Category | CR ≥ 95%<br>* | CR ≥ 90%<br>* | CR ≥ 80%<br>* | Mean MAF<br>145 ISAG<br>** | Mean MAF<br>378 Extra<br>** | Mean MAF<br>523 SNPs<br>** |
| --- | --- | --- | --- | --- | --- | --- |
| PT Samples | 507 | 523 | 537 | 0.34 | 0.40 | 0.38 |
| ISAG HCT 2019 | 541 | 548 | 555 | N/A | N/A | N/A |

\* Number of SNPs out of 562 EQ-GBS SNP panel with Call Rates $\geq 80, 90$ or $95\%$ within SNP. \*\* MAFs were calculated across 254 PT Samples, using 523 SNPs consisting of 145 pilot ISAG pilot and 378 Extra SNPs.

Duplicate genotyping of 10 ISAG HCT 2019 samples yielded a within-platform genotype concordance of 99.94% across 523 EQ-GBS SNPs. Re-genotyping of 5 of the original 40 panel design samples, yielded a 99.24% Ion S5 to Affymetrix 670K Equine Array concordance across 480 SNPs common to both genotyping panels (Table S4).

### SNP and STR parentage analysis

Pre-analysis of Mendelian SNP mismatch counts using 523 SNP panel, identified 84 Mendelian mismatch incidences spread across 26 SNPs within 55 ETP “offspring, sire and dam” trio test combinations (i.e. a Mendelian inconsistency rate of 0.29% - 84 unexpected mismatches out of a total number of 55 × 523 = 28,765 SNP site tests). Hence, the average per SNP contribution percentage of this Mendelian mismatch count was 3.85% across the 26 SNPs of which 7 SNPs exceeded this average (i.e. above 3.85%), accounting for ∼ two-thirds (57 out of 84) of observed mismatches (Table S5). These 7 SNPs were subsequently removed prior to further parentage simulation analysis, leaving a final panel of 516 EQ-GBS SNPs - inclusive of 143 ISAG pilot SNPs (Table S10).

Trio parentage combination mismatch counts for subsequent 143 and 516 SNP panels, had a maximum of 1 and 2 mismatches respectively for ETP trio combinations. ETF trio combination mismatch SNP counts displayed a normal distribution (Fig. 3) and ranged from 5 to 64 (mean 35) and 25 to 189 (mean 134) for 143 and 516 SNP panels respectively (Table 3). These findings corroborated the expected ETP and ETF cases with zero false negative or positive parentage trio combination cases for both SNP panels (Table 3 and Tables S8-S9).

**Table 3:** Marker mismatch counts for “Expected to Pass” (ETP) and “Expected to Fail” (ETF) parent combinations, separation values and number of false negative and positive cases across trio and partial sample categories for all STR and SNP datasets. Complete STR and SNP marker mismatch counts displayed in Table S6 – S9. Probability of Non-Exclusion (NE) and Exclusion (PE) are also shown for both One-Candidate-Parent (1P) and Candidate-Parent-Pair (PP) for all STR and SNP datasets.

|  |  | 12 STRs | 16 STRs | 143 SNPs | 516 SNPs |
| --- | --- | --- | --- | --- | --- |
| Trios | ETP Parent Combinations (N= 55) * | 0 - 0 (0) | 0 - 0 (0) | 0 - 1 (0.1) | 0 - 2 (0.5) |
|  | ETF Parent Combinations (N = 877,745) * | 0 - 12 (7) | 0 - 16 (9) | 5 - 64 (35) | 25 - 189 (134) |
|  | Separation Value | 0 | 0 | 0.027 | 0.044 |
|  | Number of FN cases (% out of N=55) ** | 0 | 0 | 0 | 0 |
|  | Number of FP cases (% out of N=877,745) ** | 26 (0.0030) | 2 (0.0002) | 0 | 0 |
| Partials | ETP Parent Combinations (N= 110) * | 0 - 0 (0) | 0 - 0 (0) | 0 - 1 (0.05) | 0 - 2 (0.2) |
|  | ETF Parent Combinations (N = 13,860) * | 0 - 11 (3) | 0 - 11 (4) | 1 - 28 (13) | 16 - 84 (54) |
|  | Separation Value | 0 | 0 | 0 | 0.027 |
|  | Number of FN cases (% out of N=110) ** | 0 | 0 | 0 | 0 |
|  | Number of FP cases (% out of N=13,860) ** | 223 (1.6089) | 108 (0.7575) | 1 (0.0077) | 0 (0) |
| Exclusion Probability | NE:1P (PE:1P) *** | $1.30 \times 10^{-02}$<br>(0.9870) | $5.69 \times 10^{-03}$<br>(0.9943) | $5.00 \times 10^{-07}$<br>(>0.9999) | $5.54 \times 10^{-26}$<br>(>0.9999) |
| | NE:PP (PE:PP) *** | $2.51 \times 10^{-06}$<br>(>0.9999) | $1.70 \times 10^{-07}$<br>(>0.9999) | $4.55 \times 10^{-19}$<br>(>0.9999) | $3.65 \times 10^{-70}$<br>(>0.9999) |

\* Range and Mean number of mismatch counts based on a number (N) of trio or partial parent combinations.
\*\* Exclusion threshold based on a $\geq 1\%$ SNP or $\geq 1$ STR mismatch count.
\*\*\* Estimates based on the 254 PT sample set (i.e. 309 PT samples with offspring removed).

**Fig. 3:**
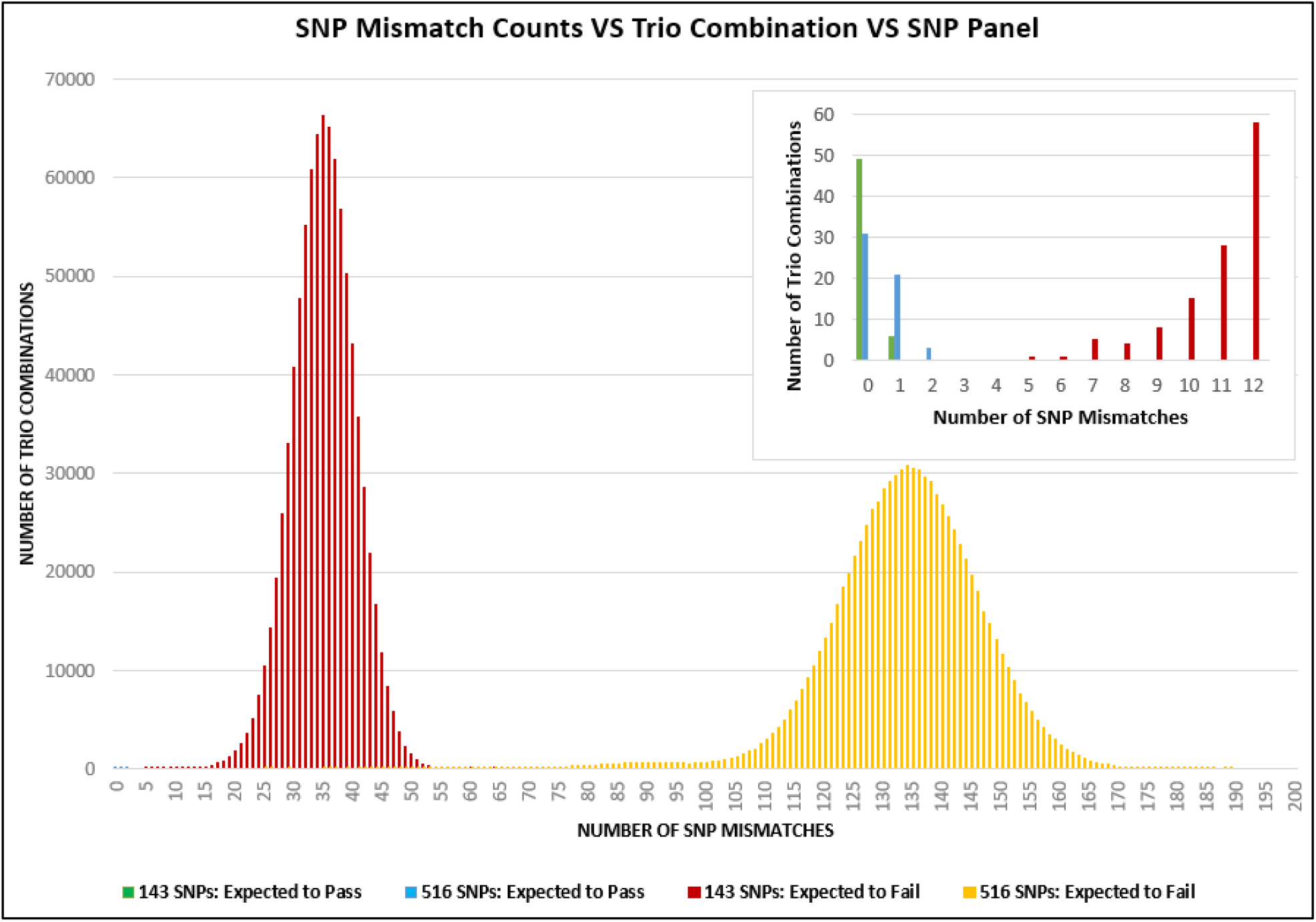
Distribution of mismatch counts for 55 “Expected to Pass” and 877,745 “Expected to Fail” simulated trio-parentage case scenarios, for both 143 and 516 SNP panels.

Trio combinations mismatch counts for both 12 and 16 STR panels were zero for ETP combinations, while mismatch counting for ETF combinations established a zero separation value for both STR panels and identified 26 (12 STR panel) and 2 (16 STR panel) false positive cases (i.e. “Expected to Fail”, however had zero mismatching STRs) out of a total of 877,745 simulated trio combinations per STR panel (Table 3). A normal mismatch count distribution was also observed for trio combination STR panels, with clear encroachment for 26 false positive cases into zero mismatch marker category for the 12 STR panel (Fig. 4).

**Fig. 4:**
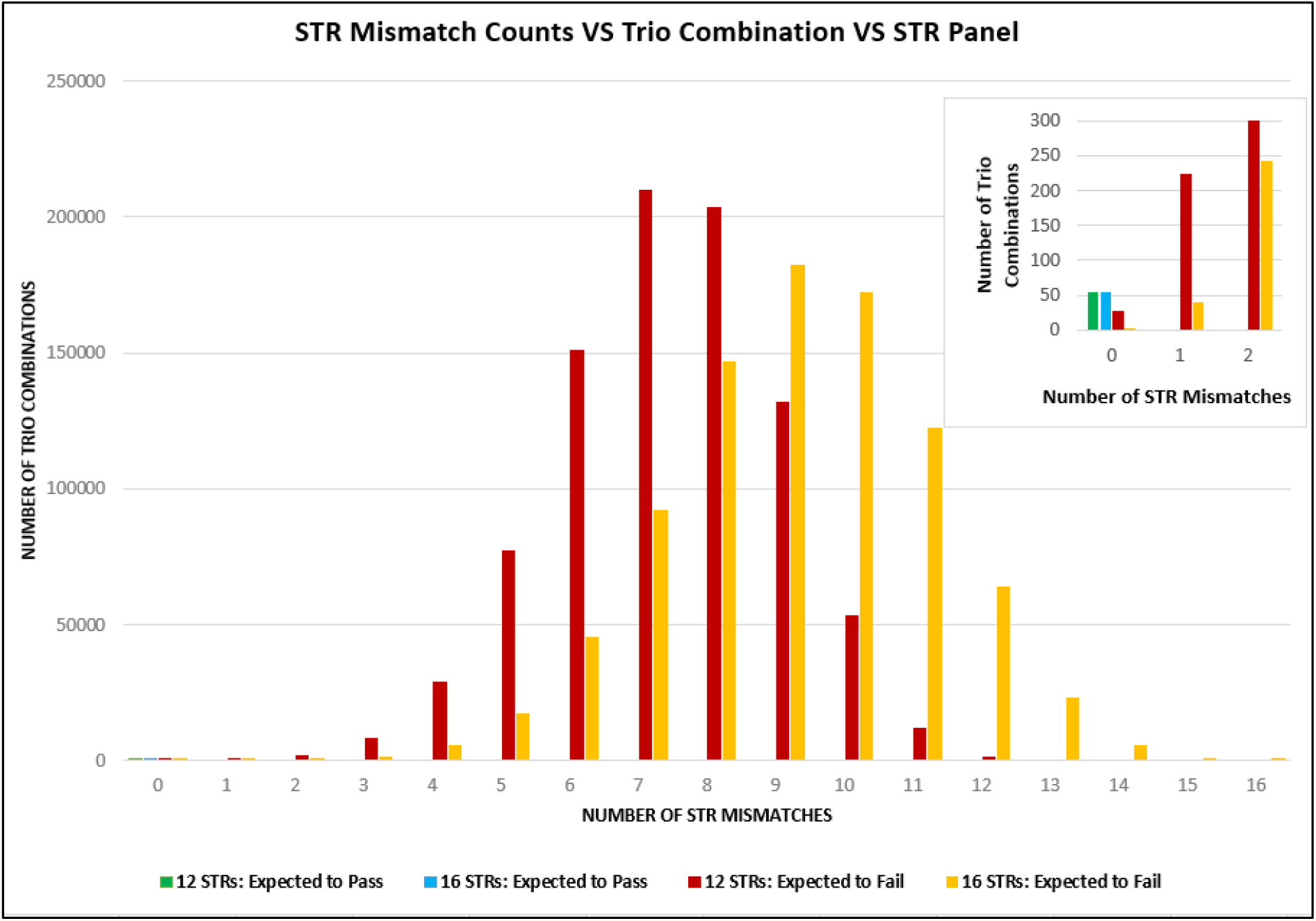
Distribution of mismatch counts for 55 “Expected to Pass” and 877,745 “Expected to Fail” simulated trio-parentage case scenarios, for both 12 and 16 STR panels.

Partial combination mismatch counts for 143 SNP, 516 SNP, 12 STR and 16 STR panels had a maximum of 1, 2, 0 and 0 mismatch counts respectively for ETP combinations. The 516 SNP panel had a minimum of 16 mismatch counts for ETF combinations, along with a positive separation value (0.027) and zero false negative/positive cases. A separation value of zero along with 1, 223 and 108 false positive cases were observed within partial combination for 143 SNP, 12 STR and 16 STR panels respectively (Table 3). A similar normal distribution mismatch count was observed for partial as with trio combinations, however a greater level of false positive encroachment was observed with 223 and 108 cases identified for 12 and 16 STR panels respectively (Fig. S6 and S7).

Probabilities of Exclusion (PE) were estimated at 0.9870 and 0.9943 for 12 and 16 STR panels, respectively, for One-Candidate-Parent (1P) category, while all remaining SNP based panel types had PEs of > 0.9999 across both One-Candidate-Parent (1P) and Candidate-Parent-Pair (PP) categories. Non-Exclusion probabilities (NE) ranged from 3.65×10^−70^ (NE:PP for 516 SNPs), to 1.30×10^−02^ (NE:1P for 12 STRs) and NE values decreased across 12 STR - 16 STR - 143 SNP - 516 SNP panels for both 1P and PP parent categories (Table 3).

## Discussion

This study demonstrates GBS as a proficient technology to generate SNP genotype profiles suitable for equine parental analysis and enhanced testing accuracies were observed when using SNPs in comparison to STRs within parentage verification and allocation scenarios.

A 98.1% synthesis conversion rate from 573 SNPs submitted for AgriSeq™ GBS panel manufacturing, yielded a 562 EQ-GBS SNP panel consisting of 151 out of 154 ISAG pilot SNPs, an extra 410 SNPs to aid parentage discovery and 1 Y-Chromosome SNP to facilitate sex quality control checks within genotyping/parentage processes. A uniform genome-wide SNP distribution (average distance between SNPs of ∼3.95Mb) was driven by both the 151 base ISAG pilot SNPs and the within chromosome linkage disequilibrium pruning selection approach taken to establish an Extra panel of 410 SNPs. Subsequent genotyping of 309 Thoroughbred horses using the 562 EQ-GBS SNP panel, identified 523 suitable SNPs i.e. average within SNP call rate of ≥90%. Furthermore, removal of 7 SNPs that had a greater than average Mendelian mismatch count (>3.85%) for true parent combinations - left a final EQ-GBS panel of 516 SNPs. All autosomal chromosomes were covered in terms of SNP representation (average ∼18.5 SNPs per chromosome) and a SNP coverage gap on ECA21, as per the 154 ISAG pilot SNP list, was attended to via addition of 12 extra SNPs located across this chromosome.

An average SNP call rate of 97.2%, across platform concordance of 99.24% and subsequent successful use of SNP profiles within parentage verification, demonstrated the EQ-GBS SNP panel’s capability to generate accurate genotype calls when using Chelex grade DNA extracts. EQ-GBS SNP genotyping of 20 ISAG HCT 2019 DNA extracts, which were extracted using commercial grade DNA extraction kits to facilitate accurate global ring trial genotyping, yielded an average SNP call rate of 99.1% i.e. ∼2 percentage units greater than when using Chelex based DNA extracts. Such a performance advantage is clearly attributable to using higher grade DNA extracts, however demonstrated feasibility of using Chelex based DNA extracts can potentially help with yet-to-be established EQ-GBS SNP genotyping price points - which is a key factor to facilitate an STR to SNP transition.

An EQ-GBS panel average MAF of 0.38 across all 523 SNPs and a within SNP call rate >90%, is comparable to previous observations for the Thoroughbred breed (Holl *et al*. 2017 and Hirota *et al*. 2010) and a multi breed MAF average of 0.33 for 371 Extra SNPs provides initial indications of the panels capability to effectively aid identity, parentage verification and parent discovery outside of other breeds than Thoroughbred. Such MAF estimates (i.e. >0.30) bode well in terms of potential parentage analysis capability (Holl *et al*. 2017) and are further bolstered due to the additional SNP numbers within this panel outside of the 154 ISAG Pilot SNP panel. Cognisance must be given to ensure this GBS panel performance holds true across equine breeds and genotyping platforms. As our multibreed MAF estimates are *in-silico* based only, with an average of ∼21 animals per breed, actual genotyping of larger sample numbers from breed-representative horses is required, for Extra SNPs within EQ-GBS panel.

Furthermore, probabilities of exclusion of >0.9999 for both PE:1P and PE:PP support the potential effectiveness of EQ-GBS SNP panel within parentage verification testing and such PE analysis provided first insights into direct statistical gain (i.e. same samples used for STR/SNP comparisons) when using SNPs in comparison to STRs – particularly for one candidate parent cases. Probability of exclusions (PE) of 0.9870 and 0.9943 were observed for PE:1P case scenarios when using 12 STR and 16 STR panels respectively, which were less than all other case/panel PE’s of >0.9999. These PE:1P reductions for 12 and 16 STRs, were borne out in terms of 1.61% and 0.76% false positives along with zero separation values when performing simulated partial parentage testing. Further false positives were also observed for both STR panels within trio parentage testing, albeit to a lesser extent of <0.01%. In general SNP panels fared better than STRs within simulated parentage analysis with positive separation values for 3 out 4 trio/partial/SNP panel scenarios, in comparison to no positive separation values for 4 trio/partial/STR panel scenarios. The only case/SNP panel scenario that displayed a separation value of 0 was the 143 SNP panel for partial testing and even though encroachment was observed for “Expected to Fail” upon “Expected to Pass” parentage combinations - only 1 false positive case out of 13,860 (0.0077%) possible case combinations was detected, regardless of there being no clear separation value. In practical terms such a false positive rate appears of little concern; however within a no prior nominated parents database search scenario, where more animal profiles may exist i.e. more potential allocation candidates, having 516 SNPs will be of benefit as demonstrated within simulated parentage testing.

Trio case scenarios are of core interest when testing Thoroughbred parentage as typically both proposed sire and dam of offspring are known. Similarly, to partial case scenarios an increase in trio case testing robustness was observed when using SNPs in comparison to STRs. A zero-separation value and a low percentage level (<0.01%) of false positives were observed for both STR panels, in comparison to positive separation values and no false positives for both SNP panels. Positive separation values for both SNP panels allowed for clear discrimination between both “Expected to Fail” and “Expected to Pass” parentage groups, and when using the 516 SNP panel a near absolute level of discrimination was observed between both combination categories (Fig. 3). This capability to establish a near absolute end point parentage result is of merit to the 516 SNP panel, and albeit the 143 SNP panel did achieve positive separation values, “Expected to Fail” parentage combination cases with ∼5 or 6 SNP mismatches can be clarified in confidence using the complete 516 SNP panel. These findings are aligned with recent studies within sheep and cattle where at least 200 SNP markers were recommended for parentage testing if the aim is to reduce false-negatives and to fully exclude false positives at least 500-700 markers are recommended (McClure *et al*. 2015 and Strucken *et al*. 2016). Furthermore, the potential value of an equine secondary SNP panel has been previously raised to address false positives/doubtful non-parent pairs when potential parents are close relatives (Holl *et al*. 2017) and simulated parentage testing using the 516 EQ-GBS SNP panel demonstrates a capability to alleviate such parentage testing concerns.

## Conclusion

SNP based parentage verification is commonly used within species such as bovine, ovine and porcine and ensuring feasibility and testing accuracy are key considerations for a potential STR to SNP transition within the equine species. Establishing a GBS based SNP panel of ∼500 SNPs demonstrated direct statistical gain and practical testing observations when using SNPs over STRs to perform parentage verification within the Thoroughbred equine breed. Genotyping further horses is required to further validate performance for the Extra SNP panel component of this EQ-GBS panel across breeds, and future panel versions will be benefited by reviewing core ISAG pilot SNP dropout along with the addition of markers to facilitate in-tandem profiling of diagnostic traits of interest to equine breeds.

Overall, benefits were observed when using 143 and 516 SNP panels for parentage testing over 12 and 16 STR panels, and recent developments within genotyping technologies such as GBS facilitate accurate generation of low-density SNP profiles providing a feasible first steppingstone to support an STR to SNP transition for parentage verification within equine species.

## Statement of ethics

No ethical approval was required as samples within this study were collected for previous parentage verification purposes, which was performed by Weatherbys Scientific.

